# A longitudinal two-photon imaging platform for focal astrocyte ablation in vivo

**DOI:** 10.64898/2026.08.26.747304

**Authors:** Nicola B. Schmid, Matthias T. Wyss, Anna Lasne, Ria Patoli, Jeffrey L. Bennett, Aiman S. Saab, Bruno Weber, Marina Herwerth

## Abstract

Investigating the consequences of astrocyte loss in the intact brain is both important and challenging. As integral components of the neuro-glia-vascular unit, astrocytes are involved in a variety of brain processes including water homeostasis, metabolic supply, regulation of cerebral blood flow, and coordination of neuronal circuit activity. Astrocyte impairment has been associated with numerous neurological disorders. However, experimental models combining focal astrocyte ablation with longitudinal *in vivo* imaging in the intact adult brain have been lacking, limiting efforts to define the causal contribution of astrocyte loss to central nervous system (CNS) pathology and repair.

Here, we present an *in vivo* model of antibody-mediated astrocyte ablation that enables longitudinal imaging and detailed investigation of ensuing cellular responses. It integrates focal induction of aquaporin-4 antibody-mediated astrocyte loss, chronic *in vivo* two-photon imaging, genetically encoded sensors, and reporter mouse lines. This advancement allows visualization and quantification of cellular and subcellular events in living organisms during lesion progression and recovery. It overcomes many longstanding limitations of previous models that are either constrained by non-specific hypoxic or mechanical tissue damage or require sacrificing animals at discrete time points, hindering the ability to monitor dynamic biological processes over time. In contrast, the selective targeting of astrocytes prevents the formation of the glial border, enabling the investigation of CNS response in a scar-free environment.

Overall, this new approach represents a significant technical advancement, enabling comprehensive longitudinal studies of CNS responses to astrocyte loss, thus opening new avenues for understanding astrocytopathy-driven pathology, evaluating therapeutic interventions, and promoting translational research.

**Significance:** Astrocyte loss has been implicated in various neurological disorders, either as a primary cause or as a contributing factor throughout disease progression. However, there remains a critical lack of experimental models capable of directly assessing the impact of astrocyte loss on CNS integrity. Here, we introduce an *in vivo* model of focal astrocyte ablation that enables multimodal, longitudinal investigation of astrocyte regeneration at both the population and single-cell levels, as well as of dynamic interactions between astrocytes and other CNS cell types. Applications of this model can substantially advance our understanding of the consequences of astrocyte loss in the adult brain, opening new opportunities to investigate its implications in neurological disorders.

## INTRODUCTION

Astrocytes are a multifunctional glial population in the central nervous system (CNS). Through their interactions with blood vessels, synapses, and microglia (1, 2), they regulate cerebral blood flow (3), preserve blood-brain barrier integrity, modulate synaptic activity, maintain ion and water homeostasis, provide a constant metabolic supply, and facilitate the clearance of metabolic waste (4–6). Despite the well-established physiological functions of astrocytes, the consequences of their selective loss for CNS integrity remain poorly understood because experimental models enabling selective and controlled astrocyte depletion have been lacking.

Several strategies have been developed to experimentally deplete astrocytes. Early studies relied on pharmacological gliotoxins, such as L-α-aminoadipate, fluoroacetate, or fluorocitrate, to impair or eliminate astrocytes (7–10). However, these compounds have limited cellular specificity, often achieve only partial depletion of astrocytes, and can directly affect neuronal metabolism and other glial cell populations. To improve selectivity, genetic ablation approaches were introduced, most commonly based on astrocyte-specific expression of herpes simplex virus thymidine kinase (HSV-TK) or inducible diphtheria toxin A (DTA), enabling targeted cell elimination following administration of the corresponding toxin (11, 12). While these models provide greater cell-type specificity, the depletion is typically widespread and difficult to restrict to defined anatomical regions, and toxin-dependent approaches can produce off-target effects. Photochemical methods such as 2Phatal offer superior spatial precision and permit cell-type-specific ablation. However, they were primarily designed for targeting single cells and are therefore not readily scalable to larger astrocyte populations (13). More recently, optogenetic and chemogenetic approaches have enabled reversible, temporally controlled manipulation of astrocyte activity, but they modulate astrocyte function rather than induce actual cell loss (14, 15). Collectively, studies using these experimental paradigms have provided important insights into astrocyte biology, yet no existing approach simultaneously achieves efficient astrocyte depletion, precise spatial and temporal control, and minimal secondary tissue responses. Consequently, distinguishing the direct consequences of astrocyte loss from secondary tissue remodeling remains a major challenge.

An expanding repertoire of human autoantibodies targeting various CNS antigens has been identified and is increasingly used to generate disease-relevant animal models (16). Yet not all of them have proven suitable for use in rodents, owing to factors such as limited antigen homology, differences in binding properties, restricted access to relevant CNS regions, and insufficient immunological responses (17). The human aquaporin-4 (AQP4), a water channel highly enriched at astrocytic endfeet in the CNS (18), shares over 99% sequence homology with its murine and rat counterparts, making it a highly attractive target for translational animal studies (19). The AQP4-specific autoantibodies (AQP4-IgG) are a pathogenic hallmark of neuromyelitis optica spectrum disorder (NMOSD) – a chronic inflammatory disease of CNS. We have previously generated bivalent IgG1 recombinant monoclonal antibodies from CSF plasma cells of an NMOSD patient (19), which reliably bind AQP4 in both cell culture and rodent tissue across multiple assay platforms. Using *ex vivo* and *in vivo* two-photon imaging of the spinal cord in mice and superfusion with an antibody/complement-containing solution, we have previously demonstrated the rapid dynamics of astrocyte depletion induced by this human-derived AQP4-IgG (20–22). However, this approach did not permit longitudinal observation, limiting our ability to further characterize astrocyte lesion progression and recovery.

Here, we developed an antibody-mediated strategy for selective, focal ablation of astrocytes *in vivo*. Recombinant human AQP4-IgG and human complement were stereotactically co-injected into the mouse somatosensory cortex, followed by implantation of a chronic cranial window for longitudinal two-photon microscopy (2PM) of cellular dynamics at both the population and single-cell levels. This method, termed the 2PM-AQP4 platform, combines efficient astrocyte depletion with precise spatial targeting and chronic optical access, enabling multimodal, longitudinal investigation of cellular and vascular dynamics in astrocyte-free cortical areas *in vivo*. Applications of the 2PM-AQP4 platform have the potential to substantially advance our understanding of CNS response to astrocyte loss in the adult brain, and to uncover novel therapeutic targets for CNS injury and disease.

## RESULTS

### Development of antibody-mediated astrocyte ablation approach for chronic two-photon imaging

A stepwise microinjection of predefined concentrations of AQP4-IgG and a complement source into the somatosensory cortex of *Aldh1l1*^GFP^ astrocyte reporter mice (Fig. 1A) generated reproducible, cylindrical astrocyte-depleted lesions with an average diameter of 200 µm and a depth of ~ 400 µm (Fig.1B-C and supplemental video 1). To minimize mechanical damage, we used pulled glass capillary pipettes with an opening diameter of 30 - 50 µm and limited the total injected volume to a maximum of 200 nanoliters (nl) distributed across the depth of the lesion. Following implantation of a chronic cranial window, astrocyte-depleted regions were repeatedly monitored *in vivo* using two-photon microscopy. In contrast, astrocyte depletion was minimal following injection of recombinant isotype-matched control-IgG together with complement (Ctrl-IgG; Fig. 1B).

**Figure 1:**
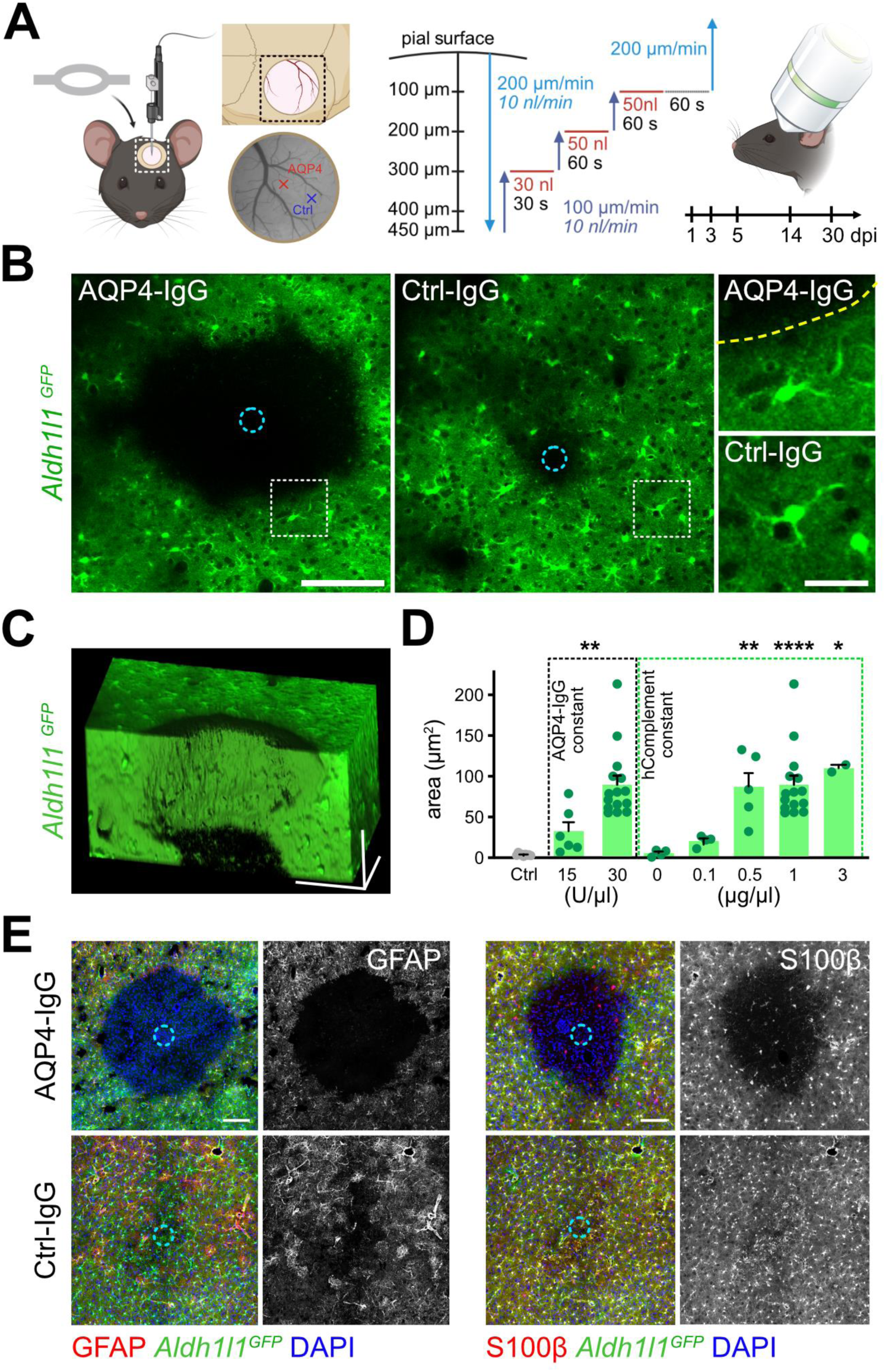
Establishment of the 2PM-AQP4 platform for focal astrocyte ablation. **A)** Scheme illustrating the injection protocol of AQP4-IgG + human complement (hC) or Ctrl-IgG + hC into the mouse cortex. **B)** One day after AQP4-IgG +hC injection (cyan dashed circles indicate injection site), focal astrocyte loss is evident in *Aldh1l1^GFP^* mice, in contrast to Ctrl-IgG + hC injected regions (scale bar = 100 µm). Right: magnification of the regions outlined by white dashed squares in the overview images (yellow dashed line marks lesion border, scale bar = 20 µm). **C)** 3D projection revealing the cylindrical shape of the lesion (scale bars in all dimensions = 100 µm). **D)** Left: quantification of depletion areas with two different complement concentrations and a fixed concentration of 1 µg/µl AQP4-IgG. (n = 6-15 animals per group; Mann-Whitney test; ** p < 0.01) Right: quantification of depletion areas across different AQP4-IgG concentrations with a fixed complement concentration (30 U/ml) (n = 2-15 animals per group; Kruskal-Wallis test; **** p < 0.0001; ** p < 0.01; * p < 0.05). Control (Ctrl): 1 µg/µl Ctrl-IgG and 30 U/ml complement (n = 5). Data are shown as mean ± SEM. **E)** Representative confocal images of horizontal sections obtained 1 dpi (scale bars = 100 µm). Sections were stained for the indicated markers in red (left: merged images, right: single channels). Endogenous *Aldh1l1^GFP^*signal shown in green and DAPI in blue in all merged images.

Next, we systematically optimized the concentrations and ratio of injected antibody and complement required for efficient astrocyte depletion. We first tested two complement doses while maintaining the AQP4-IgG titer at 1 µg/µL. A complement concentration of 30 U/ml yielded the most favorable balance between injection volume and the extent of astrocyte depletion (Fig. 1D). We subsequently tested increasing concentrations of AQP4-IgG while maintaining the complement concentration at 30 U/ml. Astrocyte depletion reached a plateau at 1 µg/µL AQP4-IgG, with no further increase in lesion size observed at higher antibody concentrations. (Fig. 1D). Therefore, 1 µg/µl AQP4-IgG and 30 U/ml complement were used in all subsequent experiments. These titration experiments demonstrate that the efficiency of astrocyte depletion depends on both antibody concentration and the availability of complement required to mediate antibody-dependent astrocyte lysis.

To verify that the disappearance of the GFP fluorescence reflected genuine astrocyte loss, mice were perfused at 1 day post-injection (dpi), brains were post-fixed, and lesion-containing sections were processed for immunohistochemical analysis. Staining for the astrocyte markers GFAP and S100β demonstrated a near-complete absence of astrocytes within the lesion core, validating efficient astrocyte ablation and excluding changes in reporter expression or imaging artifacts as explanations for the observed loss of fluorescence (Fig. 1E).

### The 2PM-AQP4 platform enables longitudinal single-cell analysis of astrocyte remodeling *in vivo*

To demonstrate the ability of the 2PM-AQP4 platform to resolve and longitudinally track structural remodeling of individual astrocytes *in vivo*, we developed a dual-labeling strategy that enables repeated visualization of individual astrocytes within the astrocyte network (Fig. 2A). Sparse astrocyte labeling was achieved in the *Glast^CreERT2^* × *Rosa26^tdTomato^* mice by administrating a single low dose of tamoxifen (50 mg/kg BW), resulting in the expression of the red fluorophore tdTomato in a subset of astrocytes. To simultaneously visualize the entire astrocyte network and accurately delineate lesion boundaries, mice additionally received an intravenous injection of the viral construct AAV-PHP.eB2-GFAP-GFP three weeks before AQP4-IgG-mediated lesion induction (Fig. 2A-B). This dual-labeling approach enabled reliable identification and repeated tracking of individual tdTomato-positive astrocytes located at the lesion border throughout the imaging period. We next assessed whether this approach could capture structural remodeling at the single-cell level. Three-dimensional reconstructions of longitudinal acquired *in vivo* image stacks allowed changes in the morphology and territorial organization of the same perilesional astrocytes to be visualized over time (Fig.2C). As a proof of principle, comparison of astrocytes at one and five dpi revealed progressive extension of individual astrocyte territories into the astrocyte-depleted region, whereas astrocytes in control areas remained morphologically stable over the same period (Fig. 2C).

**Figure 2:**
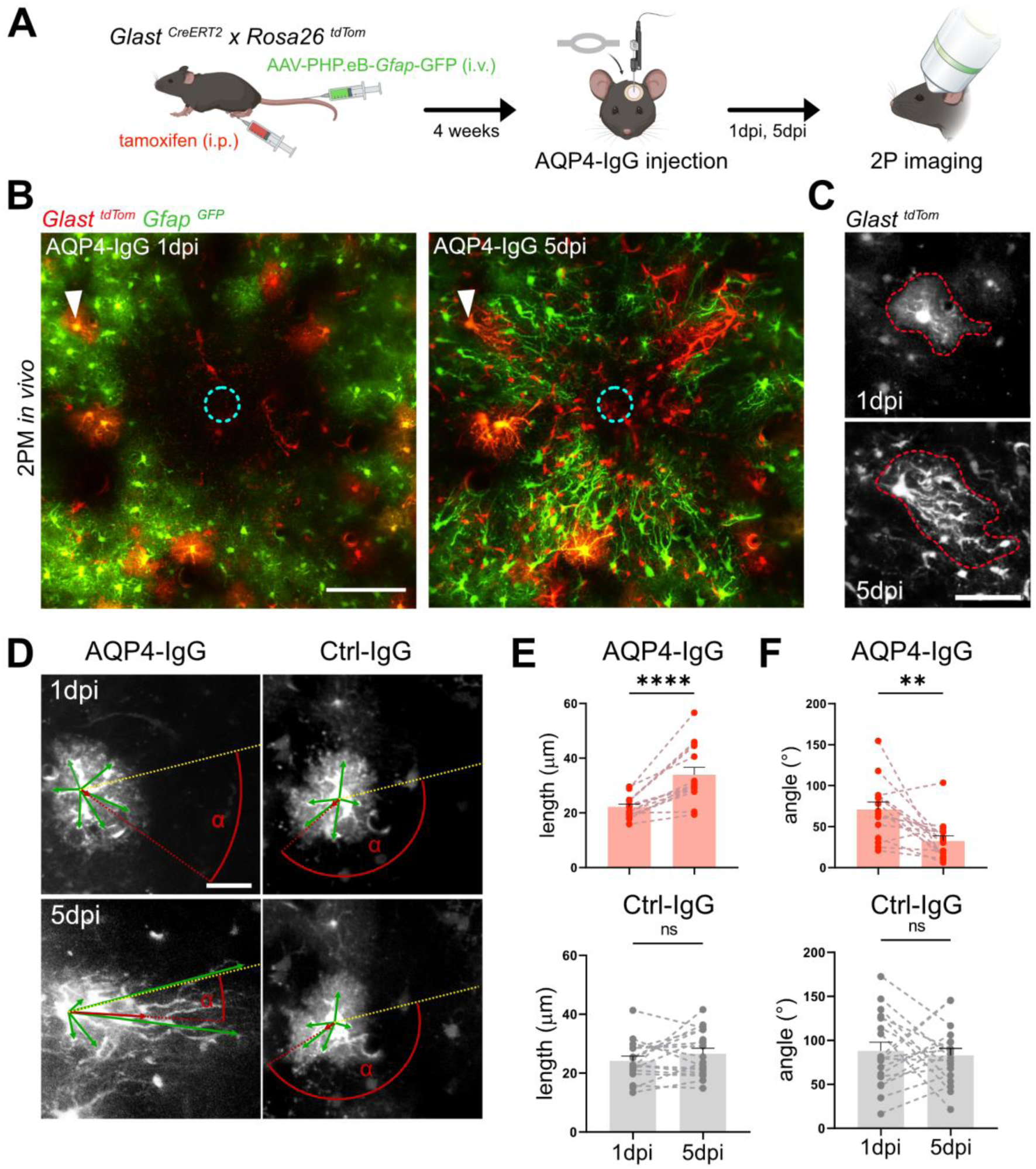
2PM-AQP4 platform allows longitudinal tracking and quantitative analysis of perilesional astrocytes. **A)** Sparse cell labeling protocol for *in vivo* chronic two-photon imaging of individual astrocytes in *Glast^CreERT2^ × Rosa26^tdTomato^* mice. **B)** Tracking single tdTomato-positive astrocytes with 2PM over time (white arrowheads, cyan dashed circle: injection site) allows detailed morphological evaluation (scale bar = 100 µm). **C)** Representative z-projection showing expansion of the territorial domain of the astrocyte marked in B between 1 and 5 dpi (scale bar = 20 µm). **D)** Representative images of individual astrocytes illustrating the vector-based analysis used to quantify astrocyte process length and orientation, angle α (red); yellow dashed lines project to the center of the lesion (COL). Green arrows show individual vectors for main astrocytic processes; the respective mean vectors are indicated by red arrows (scale bar = 20 µm) **E-F)** Vector-based quantification of astrocytic process lengths (mean vector length) (E) and polarization (F) over time (n = 15 astrocytes from 5 mice for AQP4-IgG and Ctrl-IgG, respectively, data points represent matched measurements from individual cells from 1 dpi to 5 dpi; data shown as means ± SEM. Wilcoxon signed rank test; **** p < 0.0001, ** p < 0.01, ns = not significant).

To extend the platform to quantitative analysis of cellular remodeling, we developed a three-dimensional vector-based approach to extract astrocyte process length and orientation from *in vivo* two-photon image stacks (Fig. 2D). This analysis captured both elongation of primary processes and progressive polarization toward the lesion center (Fig. 2E-F). These results demonstrate that the platform enables structural remodeling to be quantified in three dimensions and related to the spatial context of focal astrocyte loss *in vivo*.

### The 2PM-AQP4 platform provides *in vivo* optical access to the astrocyte-vascular interface

Disruption of astrocyte-vascular coupling has been implicated in numerous neuropathological conditions, including ischemic stroke, traumatic brain injury, and neurodegenerative disorders (23, 24). However, the structural complexity of the astrocyte-vascular interface is difficult to interrogate dynamically. AQP4-IgG-mediated astrocyte depletion results in the loss of perivascular astrocytic endfeet, thereby disrupting local astrocyte-vascular connectivity. We therefore asked whether the 2PM-AQP4 platform could be extended to simultaneously visualize astrocytes and the cerebral vasculature and to follow the re-establishment of astrocyte-vascular interactions following focal astrocyte depletion. For this goal, *Glast^CreERT2^ × Rosa26^tdTom^* mice were crossed with the *Claudin^GFP^* reporter line, enabling fluorescent labeling of astrocytes (red) and endothelial cells (green) within the same imaging volume following tamoxifen administration (Fig. 3A).

**Figure 3:**
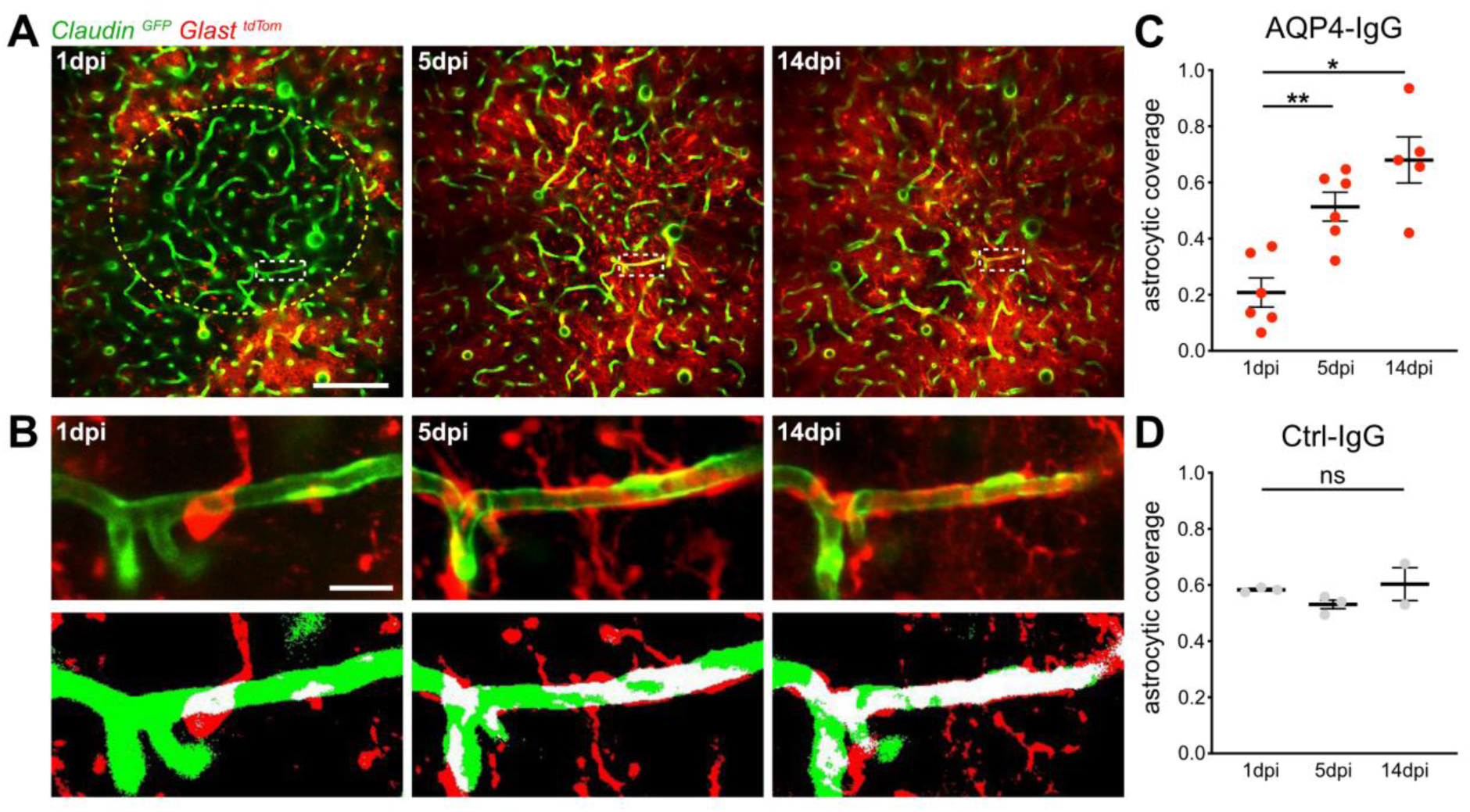
*In vivo* investigation of the astrocyte-vascular unit following astrocyte depletion. **A)** Projected two-photon image z-stacks (20 µm) from *Glast^CreERT2^ × Rosa26^tdTom^ × Claudin^GFP^* mice 1 dpi, 5 dpi, and 14 dpi (yellow dashed circle indicates lesion border, scale bar = 100 µm). **B)** Magnification from white dashed squares in (A) (upper panels) and respective segmentations (lower panels), illustrating vessel regions covered by astrocyte endfeet (white). **C-D)** Quantification of astrocytic coverage after AQP4-IgG injection (C) and after control-IgG injection (D) at different timepoints after lesion induction (AQP4-IgG: n = 6 regions from 3 animals; Ctrl-IgG: n = 3 regions from 2 animals; one-way ANOVA followed by Tukey’s multiple comparison test; * p < 0.05, ** p < 0.01, ns = non-significant; data are shown as mean ± SEM).

Chronic two-photon imaging revealed pronounced loss of astrocytic endfoot coverage one day after AQP4-IgG injection, reflecting selective disruption of astrocyte-vascular contacts within the lesion area (Fig. 3A-B). At early time points, regions with reduced astrocytic coverage of the vascular surface could be identified and repeatedly revisited. Longitudinal imaging further enabled the same vascular segments to be followed during recovery, capturing the progressive extension of astrocytic processes toward the vasculature and their re-association with the vessel surface (Fig. 3B-D). This approach further enabled quantification of vascular coverage across consecutive imaging time points, providing a longitudinal measure of structural remodeling at the astrocyte-vascular interface (Fig. 3C-D). Considering that this mouse line labels only a subpopulation of astrocytes, complete vascular coverage was not expected. Together, these findings establish the 2PM-AQP4 platform as a powerful platform for investigating the dynamics of neuro-glia-vascular remodeling following selective astrocyte loss.

### The 2PM-AQP4 platform enables analysis of astrocyte-microglia dynamics in vivo

Astrocytes and microglia engage in dynamic bidirectional crosstalk that plays a pivotal role not only in maintaining CNS homeostasis but also in orchestrating neuroinflammatory responses (25). Having established the 2PM-AQP4 platform, we next investigated whether it would enable simultaneous *in vivo* imaging and quantitative analysis of astrocyte-microglia interactions in response to astrocyte loss. To visualize both cell populations, *Glast^CreERT2^ × Rosa26^tdTom^*were crossed with *Cx3cr1^GFP^* mice, resulting in fluorescent labeling of astrocytes and microglia (Fig.4A). Longitudinal 2PM imaging following AQP4-IgG-mediated astrocyte depletion revealed dynamic morphological remodeling of perilesional microglia, whereas no such changes were observed in unaffected areas (Fig. 4A-B). The 3D *in vivo* vector-based analysis of individual perilesional microglia showed early polarization of their processes toward the lesion core, followed by progressive process retraction at 5 dpi, compared to control (Fig. 4C-D).

**Figure 4:**
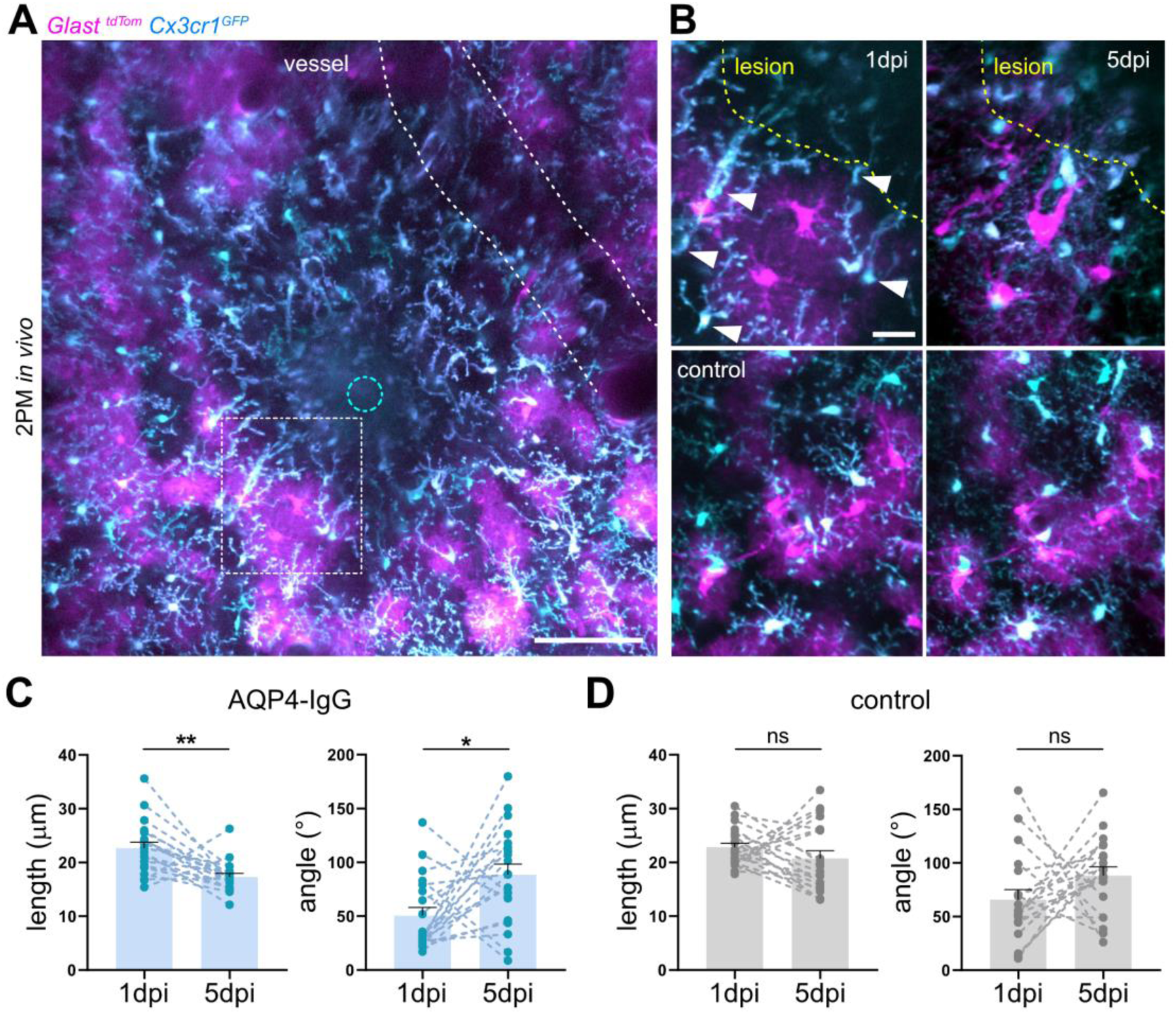
Application of the 2PM-AQP4 platform to study microglial responses to astrocyte depletion. **A)** Projected two-photon image z-stack from *Cx3cr1^GFP^ × Glast^CreERT2^ × Rosa26^tdTom^* mice, showing focal loss of astrocytes with surrounding perilesional astrocytes in magenta and microglia in cyan one day after AQP4-IgG-mediated lesion induction. Injection site is marked by a blue dashed circle (scale bar = 100 µm). **B)** Higher magnification time lapse images from the perilesional boxed area in *A* (upper panels) and a corresponding control area (lower panels), illustrating dynamic morphological changes of perilesional microglia over time (scale bar = 20 µm). White arrowheads mark polarized microglia. **C-D)** 3D vector-based quantification of microglial mean process lengths and process polarization at 1 and 5 dpi in AQP4-IgG-mediated lesions (C), and in control regions (D) (n = 20 cells from 4 mice per group; Wilcoxon signed rank test; * p < 0.05, ** p < 0.01 and ns = not significant; data are shown as mean ± SEM).

Collectively, these findings demonstrate that the 2PM-AQP4 platform is a powerful tool for longitudinal visualization and quantitative analysis of astrocyte-microglia interactions in the living brain following selective astrocyte depletion. By combining dual-color imaging with quantitative single-cell morphometric analysis, the platform establishes a versatile method for investigating glia–glia communication during tissue remodeling and neuroinflammation.

## DISCUSSION

The ability to selectively manipulate astrocytes while preserving the surrounding CNS tissue architecture remains a major challenge in the adult brain. Here, we introduce the 2PM-AQP4 platform, a two-photon-compatible model of focal AQP4-IgG-mediated astrocyte depletion that enables spatially confined and reproducible ablation of astrocytes in the intact adult cortex. When combined with chronic *in vivo* imaging, the platform provides optical access to cellular and subcellular responses following selective astrocyte loss, with high spatial and temporal resolution.

Astrocyte ablation using AQP4-targeting antibodies has previously been used and provided seminal insights into AQP4-IgG-mediated histopathology (19, 26–30). However, they were not designed for chronic high-resolution intravital imaging and often relied on endpoint histology or inflammatory disease models such as experimental autoimmune encephalomyelitis, which introduce additional systemic immune responses. As an alternative, the recently developed antibody-free 2Phatal method (13) enables selective ablation of single cells, including astrocytes, via a photochemical strategy with high spatial precision. While well-suited for single-cell studies, its scalability to larger astrocytic networks remains limited. Moreover, the kinetics of photochemically induced apoptosis (days) differ substantially from those of antibody-mediated astrocyte ablation (hours), potentially resulting in distinct cellular responses. Toxin-based strategies, such as diphtheria toxin receptor-mediated ablation, allow broader targeting but lack data supporting their use in longitudinal *in vivo* studies and offer limited control over lesion localization (11). Other models of astrocyte depletion relied on non-specific mechanical or ischemic injury, which simultaneously affect multiple cell types and therefore are not primarily targeting astrocytes, provide limited spatial control, and rapidly trigger secondary tissue damage, leading to a glial scar, thereby complicating the interpretation of astrocyte-specific effects (31–34).

Unlike previous astrocyte depletion models, the 2PM-AQP4 combines focal astrocyte depletion with chronic high-resolution imaging, allowing repeated acquisition of image volumes of up to 12 × 10^7^ µm^3^ (imaging depth 400 µm) at a 0.2-0.3 µm/pixel resolution. This enables longitudinal analysis of astrocyte responses *in vivo* and their impact on surrounding CNS structures at both single-cell and population resolution without the confounding effects of glial scar formation (35). This creates an experimental setting to dissect both the immediate microenvironmental consequences of astrocyte depletion and the subsequent regenerative processes within the CNS. In particular, the platform permits direct, real-time observation of interactions among astrocytes, other neural cell populations, and the cerebral vasculature following astrocyte loss.

As proof-of-concept applications, we demonstrate that the platform enables longitudinal quantitative analysis of astrocyte remodeling, progressive re-establishment of astrocyte–vascular interactions, and dynamic microglial responses following selective astrocyte depletion. Combined with multicolor fluorescence imaging, three-dimensional image reconstruction, and quantitative morphometric analyses, these examples illustrate the versatility of the platform for investigating cellular interactions at both single-cell and population resolution. Beyond the applications presented here, the approach is compatible with a broad range of genetically encoded reporters and biosensors, allowing the integration of structural, functional, and metabolic readouts within the same experimental preparation. Together, these features enable repeated interrogation of cellular behavior within the same tissue region over time, opening new opportunities to unravel the dynamic processes that underlie tissue remodeling following astrocyte loss.

The flexibility of the 2PM-AQP4 platform provides numerous opportunities for future applications. It is essentially compatible with any disease model or transgenic mouse line and can be readily combined with histology, spatial transcriptomics, and advanced imaging approaches to link longitudinal *in vivo* observations with molecular profiling. Likewise, genetically encoded calcium indicators, metabolic sensors, or functional paradigms such as targeted whisker stimulation can be incorporated to investigate the functional consequences of astrocyte loss on cortical circuit activity. Given the comparable sizes of an astrocyte depletion lesion with individual whisker barrels in the rodent somatosensory cortex (~200-300 µm (36), the platform would elegantly allow the investigation of consequences of missing astrocytes on the processing of the input of individual whiskers. These features broaden its applicability beyond structural analyses and position the platform as a versatile tool for studying glial biology, neurovascular coupling, and neuroimmune communication *in vivo*.

Beyond its value as a general experimental platform, the model also reproduces key aspects of the initiating AQP4-IgG-mediated astrocyte loss that characterizes NMOSD, thereby providing a clinically relevant framework for investigating antibody-mediated CNS astrocytopathies (37). Although considerable progress has been made in understanding NMOSD pathogenesis (37–40), the mechanisms linking primary astrocyte loss to subsequent neuronal dysfunction and tissue degeneration remain incompletely understood. The 2PM-AQP4 platform could be particularly useful to explore the impact of astrocyte loss on neuronal circuit function *in vivo*, providing a deeper understanding of the pathological cascades underlying NMOSD.

Several limitations of the 2PM-AQP4 platform should be considered. Although AQP4-IgG-mediated complement activation produces selective astrocyte depletion, it inevitably induces a transient local neuroinflammatory milieu that must be taken into account when interpreting downstream cellular responses. Consequently, appropriate experimental controls remain essential when attributing biological effects specifically to astrocyte loss. In addition, while the model faithfully reproduces the initiating astrocytopathy characteristic of NMOSD, it does not capture the full complexity of the peripheral and adaptive immune responses that contribute to lesion formation and disease course in patients. Moreover, the recombinant patient-derived AQP4-IgG antibodies used in this study are currently not commercially available, which may limit immediate implementation of the platform by other laboratories. Finally, the broader applicability of this approach to other antibody-mediated CNS disorders may be limited. The 2PM-AQP4 platform exploits the exceptional cross-species conservation of AQP4, enabling patient-derived human AQP4-IgG to efficiently recognize the endogenous murine antigen and trigger complement-mediated astrocyte depletion. Extending this strategy to other autoimmune CNS diseases will depend on comparable antigen homology and preservation of antibody pathogenicity across species, which cannot be assumed for all disease-associated autoantibodies.

In summary, the 2PM-AQP4 platform provides a reproducible and versatile platform for investigating the consequences of selective astrocyte depletion in the intact adult brain with unprecedented spatial and temporal resolution. By combining focal antibody-mediated lesion induction with chronic high-resolution imaging and broad compatibility with fluorescent reporters, biosensors, and emerging imaging technologies, the platform enables direct visualization of cellular interactions, structural remodeling, and functional adaptations following astrocyte loss. As such, it offers a powerful methodological framework for studying astrocyte plasticity, neuro-glia-vascular interactions, and antibody-mediated neuroinflammatory disease *in vivo*.

## MATERIAL AND METHODS

### Animals

Male and female mice aged 4-6 months were used in this study. All animals were kept in temperature- and humidity-controlled husbandry conditions (22-24 °C, 50-60% relative humidity) with unlimited access to food and water. Male and female littermates were equally allocated into control and experimental groups. All animal experiments were approved by the local veterinary authorities (Veterinary Office of the Canton of Zurich) in accordance with the prescriptions and guidelines of the Swiss Animal Protection Law (Animal Welfare Act of 16 December 2005 and Animal Protection Ordinance of 23 April 2008).

To visualize astrocytes, the aldehyde dehydrogenase 1 family member L1 *(Aldh1l1)^GFP^* mouse strain was obtained from MMRRC (Tg(Aldh1l1-EGFP)OFC789Gsat/Mmucd). To trace individual astrocytes, Glutamate Aspartate Transporter (*Glast)^CreERT2^* (Strain #:012586) × *Rosa26^tdTomato^* mice (Strain #:007914) were injected once at a low dose of tamoxifen (50 mg/kg intraperitoneally (i.p.), 5 mg/ml in corn oil) four weeks before imaging to obtain sparse red fluorescent astrocyte labeling. Additionally, ssAAV-PHP.eB/2-hGFAP-EGFP-WPRE-hGHp(A) virus (50 µl of 2.0 × 10^13^ particles/ml, Viral Vector Facility, UZH, Zurich, Switzerland) was injected intravenously (i.v.) four weeks before imaging to obtain widespread astrocytic GFP expression for identification of the lesion area.

In experiments investigating the interaction of astrocytic endfoot interaction with brain vasculature, Glast*^CreERT2^* (Strain #:012586) × *Rosa26^tdTomato^* mice (Strain #:007914) were obtained from Jackson Laboratory and crossed with the *Claudin^GFP^* mouse strain (B6.Cg-Tg (Cldn5-EGFP)Cbet/U) to achieve simultaneous labeling of vasculature (green) and astrocytes (red). Tamoxifen was injected once (100 mg/kg i.p., 10 mg/ml in corn oil) four weeks before imaging to obtain Cre recombinase and tdTomato expression.

To visualize astrocytes and microglia simultaneously, *Glast^CreERT2^* (Strain #:012586) × *Rosa26^tdTomato^* mice (Strain #:007914) were crossed with the *Cx3cr1^GFP^* (Strain: #005582) mouse line. Tamoxifen was injected once (100 mg/kg i.p., 10 mg/ml in corn oil) four weeks before imaging to obtain Cre recombinase and tdTomato expression.

### AQP4 and isotype control antibodies

A detailed protocol for the generation of AQP4-IgG has previously been published elsewhere (19). In brief, human IgG1 recombinant AQP4-IgG (clone 7-5-53) was reconstructed from a clonotypic plasma blast, which was obtained from the cerebrospinal fluid of an NMOSD patient. Control-IgG of the same IgG subtype as AQP4-IgG (human IgG1) of unknown specificity was gained from a meningitis patient (clone 2B4).

### Surgical interventions

For surgical interventions, animals were anesthetized by i.p. injection of a mixture of fentanyl (0.05 mg/kg bodyweight (BW); Sintenyl; Sintetica), midazolam (5 mg/kg BW, Roche), and medetomidine (0.5 mg/kg BW, Orion Pharma) which was reapplied as needed or after 60 minutes. During anesthesia, vitamin A eye ointment (Vitamin A, Bausch+Lomb) was applied to prevent corneal desiccation. To prevent dehydration, 10 ml/kg of prewarmed Ringerfundin was subcutaneously (s.c.) injected before the intervention. Oxygen was delivered (200 cc/min) to prevent hypoxemia. Animals’ vital signs and body temperature were monitored via MARTA Pad (Vigilitech).

For head plate implantation, animals were fixed in a stereotaxic frame (Model 900; David Kopf Instruments). The head region was shaved, disinfected (Kodan; Schülke & Mayr), and anesthetized by local s.c. injection of a mixture of lidocaine (10 mg/ml, Streuli Pharma) and bupivacaine (5 mg/ml, Syntetica). The skull surface was exposed by removing a circular patch of skin above the planned craniotomy (1 - 1.5 cm). Connective tissue was removed and the skull was covered with a bonding agent (One Coat 7 Universal, Coltene). Next, a custom-made stainless steel head plate was positioned centrally above the exposed bone and secured in place by multiple layers of blue light-curing dental cement (Tetric EvoFlow, Ivoclar Vivodent), while sparing the area designated for later craniotomy.

A craniotomy above the left somatosensory cortex was performed using a dental drill (diameter 0.2 mm, H-4-002HP, Rotatec GmbH). The dura was removed, and a mixture of AQP4-IgG (0.1 - 3 µg/µl) or Control-IgG (0.1 - 3 µg/µl) together with human complement (15-30 U/ml, Sigma-Aldrich, #S1764) was injected using a custom-made microinjector. The mixture was injected through a pulled glass capillary pipette (Drummond PCR micropipettes, Drummond Scientific) with a tip diameter of 30-50 µm. A volume of 130 nl was distributed between 0-300 µm depth in 100 µm increments and an additional 60 nl were slowly infused (10 nl/min) during the insertion and retraction of the pipette to maintain its permeance. In control lesions, Ctrl-IgG (0.1-3 µg/µl, clone 2B4) was injected together with complement in the same fashion. For chronic 2PM measurements, a round glass window (3.5 mm diameter) was implanted after completion of the antibody injection. The glass was gently placed and sealed with light-curing dental cement (Tetric Evoflow, Ivoclar Vivodent). Anesthesia was reversed by i.p. injection of a mixture of flumazenil (0.9mg/kg BW, Cito Pharma) and atipamezole (45 mg/kg BW, Virbac). After surgeries, animals were treated with buprenorphine (0.1 mg/kg s.c., Streuli) and carprofen (10 mg/kg s.c., Zoetis). Post-operative analgesia with carprofen was maintained for at least three days.

### Anesthetized *in vivo* two-photon microscopy imaging

Two-photon imaging was performed with a custom-made two-photon laser scanning microscope (41) equipped with two two-photon lasers (Chameleon Discovery NX, Coherent) and a 25x water-immersion objective (W Plan-Apochromat 25x/1.05 NA, Olympus). The tunable laser was set to 920 nm to excite GFP. The second laser with a fixed 1040 nm wavelength was used to excite tdTomato. The emitted light was collected by photomultiplier tubes (H9305-03, Hamamatsu) equipped with emission filters for blue fluorescence (475/64 Brightline HC; AHF Analysentechnik), green (535/50 Brightline HC, AHF Analysentechnik), and red fluorescence (607/70, 607/70 Brightline HC, AHF Analysentechnik). A dichroic mirror at 506 nm and 560 nm separated the excitation and emission light. ScanImage (42) and custom-written LabVIEW software were used for image control and data acquisition. Mice were anesthetized with isoflurane (1.5 - 2% in O_2_ and air). Twenty-four hours after antibody injection, the first images were acquired followed by predefined time points (see results). Stacks were acquired with a resolution of 512 × 512 pixels at 0.74 Hz and 5 μm step size for overview and with a resolution of 2048 × 2048 pixels at 0.37 Hz and 2 μm step size for detailed signal analysis.

### Image analysis

Images were processed using the open-source image analysis software FIJI (ImageJ 2.1.0) (43). For display purposes, two-dimensional (2D) images were generated using z-maximum intensity projections. In non-quantitative panels, the gamma value was adjusted non-linearly to enhance the visibility of low-intensity objects. Datasets were processed with Excel (Microsoft Corporation), and graphs were generated using Prism 9 (Graphpad). All figures were crafted with Affinity Designer 2 (Serif). Graphical illustrations of experimental protocols were generated with BioRender.

#### Lesion size analysis

*In vivo* image stacks of *Aldh1l1^GFP^*mice were z-projected over 50-150 µm depth and used for lesion area calculation.

#### Vector analysis

Vector analysis was performed as previously described (35). In brief, astrocytes or microglia were randomly selected from perilesional areas. Distal ends of primary processes, location of cell soma, as well as the centre of the lesion, were manually marked by single-point ROIs in two-photon image stacks taken at different time points after AQP4-IgG-mediated lesion induction. Coordinates of ROIs in x, y, and z were extracted. From these coordinates, every process was assigned a vector that reflected its length and orientation. This allowed for the calculation of an average vector for every cell (see Equation 1) as well as average process length and process number by means of a custom-written Python script (Python 3.6.5), summarising both length and orientation of the cell’s primary processes. In controls, astrocytes, located at the average radius of AQP4-IgG-mediated lesion 1 dpi were randomly assigned for analysis. Microglial morphology was compared to randomly chosen cells in unaffected adjacent areas.

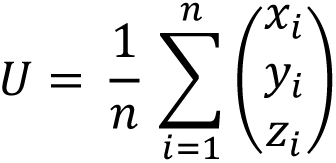

**Equation 1**: Calculation of the coordinates of the average vector U by averaging x, y, and z coordinates of all vectors of a cell.

#### Vessel coverage analysis

ImageJ was used to determine the extent of vascular regions that are occupied by astrocytic endfeet. Acquired z-stacks were split into separate channels. Thresholds were applied to segment vessels and astrocytes respectively, and binary images were created. Using Image Calculator (Operation: AND) from the two masks, a resulting image displaying vessel regions covered by astrocytic endfeet was calculated. Area measurements were applied to obtain the total vessel area and the area covered by astrocytic endfeet, from which endfoot coverage (%) was calculated (overlap area/vessel area × 100).

### Immunohistochemistry and confocal microscopy

Animals were i.p. injected with pentobarbital (200 mg/kg BW, Kantonsapotheke Zürich) for terminal anaesthesia. Animals were perfused transcardially with 4% paraformaldehyde (PFA) in 0.01 M phosphate-buffered saline (1 × PBS; in mM: 1.5 KH_2_PO_4_, 2.7 KCl, 8.1 Na_2_HPO_4_, and 137 NaCl). Brains were dissected, postfixed for 3 hours in 4% PFA, and transferred to 30% sucrose (Sigma) solution in PBS for cryoprotection overnight at 8°C. Brains were placed in embedding medium (NEG-50®, Thermo Fisher Scientific) with the apical cortical surface facing the cutting plane and then cut into 30 μm sections in a cryostat (LEICA CM3050S, Leica Biosystems AG). Antibodies were diluted in 0.3% Triton X-100 (Sigma) and 5% normal donkey serum (ab7475, Abcam) in Tris-buffered saline. The following antibodies were used: anti-GFAP (1:500-700, chicken, ab4674, Abcam), anti-S100β (1:700, rabbit, 287004, Synaptic Systems). Slices were incubated with the appropriate secondary antibodies and the nuclear dye 4’, 6-diamidino-2-phenylindole (DAPI, 1:10’000, ab228549, Abcam). Slides were imaged with a Zeiss confocal microscope (CLSM 800) equipped with ZEN 2011 (black edition, V7.1) using a 10x air objective (Plan-Apochromat 10x/0.45 M27, Zeiss) and 25x oil-immersed objective (LCI Plan-Neofluar 25x/0.8 Imm Corr DIC M27, Zeiss).

### Statistical analysis

Statistical analysis was performed in GraphPad Prism 9 software. Data is presented as mean ± SEM. All datasets were tested for normality using Shapiro-Wilk test. If the normality test was passed (p < 0.05), parametric tests were chosen. Otherwise, nonparametric tests were applied. For normally distributed data, two-tailed paired t-test was used to compare two paired groups and one-way ANOVA followed by Tukey’s multiple comparison test was used to compare multiple groups. For non-normally distributed data, statistical significance was calculated using the Mann-Whitney test when comparing two unpaired groups, the Kruskal-Wallis test followed by Dunn’s multiple comparison test when comparing more than two groups, and Wilcoxon signed-rank test when comparing two paired groups. P-values < 0.05 were considered statistically significant and indicated with asterisks in graphs as: * p < 0.05, ** p < 0.01, *** p < 0.001, and p **** < 0.0001.

## ACKNOWLEDGEMENTS AND FUNDING SOURCES

We thank S. Weber, H. Osswald, N. Binini, and A. Siebert for technical and administrative support. We thank Jean-Charles Paterna and the Viral Vector Facility of the Neuroscience Center Zurich (ZNZ) for viral productions. The *Aldh1l1*^GFP^ mouse strain [STOCK Tg(Aldh1l1-EGFP)OFC789Gsat/Mmucd; identification no.: 011015-UCD] was obtained from the Mutant Mouse Regional Resource Center, a NCRR-NIH-funded strain repository, and was donated to the MMRRC by the NINDS funded GENSAT BAC transgenic project. We sincerely thank C. Betsholtz for the CLDN5-GFP mice.

MH was supported by the Deutsche Forschungsgemeinschaft (DFG) research grant (#444138499), by the UZH Candoc Postdoc Grant (#FK-22-048), the ZNZ PhD Grant (2025), the Swiss MS Society Grant and the Swiss National Science Foundation (SNSF) Ambizione Grant (PZ00’3_216616/1). MTW was supported by the Stiftung für wissenschaftliche Forschung UZH (STWF-22-013). ASS was supported by the SNSF (Eccellenza 187000). JLB was supported by the National Institute of Neurological Disorders and Stroke (R01NS130100). BW was supported by the SNSF (31003A_156965).

## COMPETING INTEREST DECLARATION

MH received speaker honoraria, travel support and/or served on scientific advisory boards of Biogen, Sanofi, Merck Serono, Alexion, Horizon Therapeutics (Amgen), Neuraxpharm and Roche. Her institution received a research grant from Roche. JLB reports personal fees from AbbVie, Alexion, Antigenomycs, BeiOne, Chugai, Clene Nanomedicine, EMD Serono, Genentech, Genzyme, Horizon Therapeutics, Mitsubishi Tanabe Pharma, MedImmune/Viela Bio, Novartis, Reistone Biopharma, Roche, and TG Therapeutics for consultative work and scientific advisory boards; grants from Novartis, Mallinckrodt, and Alexion, and a patent for aquaporumab. None of these activities present a conflict of interest relevant to the study’s topic.

## Author contributions

NBS, MTW, BW, and MH are responsible for the concept and study design. NBS, MTW, AL, JC, RP, LR, JLB, ASS, and MH were involved in sample/data acquisition and analysis. NBS, MTW, BW, and MH drafted the manuscript and figures with input from all authors.

## Competing Interest Statement

nothing to declare

## Notes

### Competing Interest Statement

The authors have declared no competing interest.

## REFERENCES

1. L. F. Barros, S. Schirmeier, B. Weber, The Astrocyte: Metabolic Hub of the Brain. Cold Spring Harb Perspect Biol 16 (2024).

2. E. C. Damisah et al., Astrocytes and microglia play orchestrated roles and respect phagocytic territories during neuronal corpse removal in vivo. Sci Adv 6, eaba3239 (2020).

3. T. Takano et al., Astrocyte-mediated control of cerebral blood flow. Nat Neurosci 9, 260–267 (2006).

4. R. D. Fields, B. Stevens-Graham, New insights into neuron-glia communication. Science 298, 556–562 (2002).

5. A. Volkenhoff et al., Glial Glycolysis Is Essential for Neuronal Survival in Drosophila. Cell Metab 22, 437–447 (2015).

6. B. Weber, L. F. Barros, The Astrocyte: Powerhouse and Recycling Center. Cold Spring Harb Perspect Biol 7 (2015).

7. R. A. Swanson, S. H. Graham, Fluorocitrate and fluoroacetate effects on astrocyte metabolism in vitro. Brain Res 664, 94–100 (1994).

8. D. R. Brown, H. A. Kretzschmar, The glio-toxic mechanism of alpha-aminoadipic acid on cultured astrocytes. J Neurocytol 27, 109–118 (1998).

9. M. F. Pereira et al., l-alpha-aminoadipate causes astrocyte pathology with negative impact on mouse hippocampal synaptic plasticity and memory. FASEB J 35, e21726 (2021).

10. H. Zhuang et al., The Dose-Dependent Effects of Fluorocitrate on the Metabolism and Activity of Astrocytes and Neurons. Brain Sci 15 (2025).

11. B. Schreiner et al., Astrocyte Depletion Impairs Redox Homeostasis and Triggers Neuronal Loss in the Adult CNS. Cell Rep 12, 1377–1384 (2015).

12. R. R. Voskuhl et al., Reactive astrocytes form scar-like perivascular barriers to leukocytes during adaptive immune inflammation of the CNS. J Neurosci 29, 11511–11522 (2009).

13. R. A. Hill, E. C. Damisah, F. Chen, A. C. Kwan, J. Grutzendler, Targeted two-photon chemical apoptotic ablation of defined cell types in vivo. Nat Commun 8, 15837 (2017).

14. J. Bang, H. Y. Kim, H. Lee, Optogenetic and Chemogenetic Approaches for Studying Astrocytes and Gliotransmitters. Exp Neurobiol 25, 205–221 (2016).

15. X. Yu, J. Nagai, B. S. Khakh, Improved tools to study astrocytes. Nat Rev Neurosci 21, 121–138 (2020).

16. E. Maudes, J. Planaguma, M. S. Weber, J. Dalmau, Animal models of autoimmune encephalitis. Curr Opin Immunol 95, 102579 (2025).

17. N. Sinmaz, T. Nguyen, F. Tea, R. C. Dale, F. Brilot, Mapping autoantigen epitopes: molecular insights into autoantibody-associated disorders of the nervous system. J Neuroinflammation 13, 219 (2016).

18. J. M. Crane, A. S. Verkman, Determinants of aquaporin-4 assembly in orthogonal arrays revealed by live-cell single-molecule fluorescence imaging. J Cell Sci 122, 813–821 (2009).

19. J. L. Bennett et al., Intrathecal pathogenic anti-aquaporin-4 antibodies in early neuromyelitis optica. Ann Neurol 66, 617–629 (2009).

20. M. Herwerth et al., In vivo imaging reveals rapid astrocyte depletion and axon damage in a model of neuromyelitis optica-related pathology. Ann Neurol 79, 794–805 (2016).

21. M. Herwerth et al., A new form of axonal pathology in a spinal model of neuromyelitis optica. Brain 145, 1726–1742 (2022).

22. S. R. Kalluri et al., P2R Inhibitors Prevent Antibody-Mediated Complement Activation in an Animal Model of Neuromyelitis Optica: P2R Inhibitors Prevent Autoantibody Injury. Neurotherapeutics 19, 1603–1616 (2022).

23. D. Attwell et al., Glial and neuronal control of brain blood flow. Nature 468, 232–243 (2010).

24. B. A. MacVicar, E. A. Newman, Astrocyte regulation of blood flow in the brain. Cold Spring Harb Perspect Biol 7 (2015).

25. A. Nimmerjahn, F. Kirchhoff, F. Helmchen, Resting microglial cells are highly dynamic surveillants of brain parenchyma in vivo. Science 308, 1314–1318 (2005).

26. C. Wrzos et al., Early loss of oligodendrocytes in human and experimental neuromyelitis optica lesions. Acta Neuropathol 127, 523–538 (2014).

27. A. Winkler et al., Blood-brain barrier resealing in neuromyelitis optica occurs independently of astrocyte regeneration. J Clin Invest 131 (2021).

28. L. Tradtrantip, X. Yao, T. Su, A. J. Smith, A. S. Verkman, Bystander mechanism for complement-initiated early oligodendrocyte injury in neuromyelitis optica. Acta Neuropathol 134, 35–44 (2017).

29. C. Qin et al., Soluble TREM2 triggers microglial dysfunction in neuromyelitis optica spectrum disorders. Brain 147, 163–176 (2024).

30. T. Chen et al., Astrocyte-microglia interaction drives evolving neuromyelitis optica lesion. J Clin Invest 130, 4025–4038 (2020).

31. F. D. Shi, Astrocyte in stroke: Location matters. Neuron 113, 4089–4091 (2025).

32. I. B. Wanner et al., Glial scar borders are formed by newly proliferated, elongated astrocytes that interact to corral inflammatory and fibrotic cells via STAT3-dependent mechanisms after spinal cord injury. J Neurosci 33, 12870–12886 (2013).

33. S. Bardehle et al., Live imaging of astrocyte responses to acute injury reveals selective juxtavascular proliferation. Nat Neurosci 16, 580–586 (2013).

34. M. A. Anderson et al., Astrocyte scar formation aids central nervous system axon regeneration. Nature 532, 195–200 (2016).

35. M. Herwerth et al., Focal astrocyte loss reveals nuclear translocation during lesion repopulation. Nat Neurosci 29, 1826–1840 (2026).

36. C. C. H. Petersen, Sensorimotor processing in the rodent barrel cortex. Nat Rev Neurosci 20, 533–546 (2019).

37. D. M. Wingerchuk, C. F. Lucchinetti, Neuromyelitis Optica Spectrum Disorder. N Engl J Med 387, 631–639 (2022).

38. C. F. Lucchinetti et al., The pathology of an autoimmune astrocytopathy: lessons learned from neuromyelitis optica. Brain Pathol 24, 83–97 (2014).

39. A. M. Afzali et al., B cells orchestrate tolerance to the neuromyelitis optica autoantigen AQP4. Nature 10.1038/s41586-024-07079-8 (2024).

40. V. A. Lennon, T. J. Kryzer, S. J. Pittock, A. S. Verkman, S. R. Hinson, IgG marker of optic-spinal multiple sclerosis binds to the aquaporin-4 water channel. J Exp Med 202, 473–477 (2005).

41. J. M. Mayrhofer et al., Design and performance of an ultra-flexible two-photon microscope for in vivo research. Biomed Opt Express 6, 4228–4237 (2015).

42. T. A. Pologruto, B. L. Sabatini, K. Svoboda, ScanImage: flexible software for operating laser scanning microscopes. Biomed Eng Online 2, 13 (2003).

43. J. Schindelin et al., Fiji: an open-source platform for biological-image analysis. Nat Methods 9, 676–682 (2012).

